# The human Shu complex promotes RAD51 activity by modulating RPA dynamics on ssDNA

**DOI:** 10.1101/2024.02.14.580393

**Authors:** Sarah R. Hengel, Katherine Oppenheimer, Chelsea Smith, Matthew A. Schaich, Hayley L. Rein, Julieta Martino, Kristie Darrah, Oluchi Ezekwenna, Kyle Burton, Bennett Van Houten, Maria Spies, Kara A. Bernstein

**Author notes:** Corresponding authors: Kara A. Bernstein, and Sarah R. Hengel, @science_SRH.

## Abstract

Templated DNA repair that occurs during homologous recombination and replication stress relies on RAD51. RAD51 activity is positively regulated by BRCA2 and the RAD51 paralogs. The Shu complex is a RAD51 paralog-containing complex consisting of SWSAP1 and SWS1. We demonstrate that SWSAP1-SWS1 binds RAD51, maintains RAD51 filament stability, and enables strand exchange. Using single molecule confocal fluorescence microscopy combined with optical tweezers, we show that SWSAP1-SWS1 decorates RAD51 filaments proficient for homologous recombination. We also find SWSAP1-SWS1 enhances RPA diffusion on ssDNA. Importantly, we show human *sgSWSAP1* and *sgSWS1* knockout cells are sensitive to pharmacological inhibition of PARP and APE1. Lastly, we identify cancer variants in SWSAP1 that alter SWS1 complex formation. Together, we show that SWSAP1-SWS1 stimulates RAD51-dependent high-fidelity repair and may be an important new cancer therapeutic target.

## Introduction

RAD51 is the main recombinase employed during homologous recombination to enable high-fidelity templated repair and resolution of replication stress^1^. RAD51 activities are regulated by a group of proteins known as the RAD51 mediator protein complexes including the BCDX2, CX3, PALB2-RAD51C-BRCA2, and the Shu complex^2–4^. However, how these RAD51 mediator complexes specifically function to promote homologous recombination and to counteract replication stress remains poorly defined. Furthermore, it is enigmatic why so many different RAD51 mediators are needed.

The Shu complex is conserved in every eukaryotic lineage and is comprised of SWS1 and SWS1-associated protein 1, SWSAP1, which contains Walker A and Walker B motifs^5–8^. We recently identified two additional interacting partners to include the RAD51-interacting protein, scaffold protein involved in DNA repair, SPIDR, and the recombination mediator, PDS5B^6,9^. SWSAP1 is a RAD51 paralog. Like RAD51, RAD51 paralogs contain Walker A and Walker B motifs, which likely coordinate ATP binding and hydrolysis. The precise function of how SWSAP1-SWS1 modulates RAD51 activities during homologous recombination and replication stress remains unknown.

Hints to the human Shu complex function stemming from our work, and others, have shown that yeast Shu complex mutants are sensitive to the alkylating agent methylmethane sulfonate (MMS) and that the yeast Shu complex promotes Rad51 pre-synaptic filament assembly^7,10,11^. Furthermore, we found that the yeast Shu complex preferentially enables bypass of specific replication fork blocking lesions, including abasic sites, by a RAD51-mediated template switching mechanism^12,13^. The function of the human Shu complex in RAD51-dependent repair has yet to be mechanistically defined.

Using a reconstituted system, we show that the human Shu complex, SWSAP1-SWS1, plays multiple novel roles regulating RAD51 functions during high-fidelity repair in the presence of RPA. We show that SWSAP1-SWS1 forms a heterodimeric protein complex in solution that is positionally decorated throughout the RAD51 filament. We show that mechanistically SWSAP1-SWS1 binds to RAD51 maintaining the ssDNA-RAD51 interaction and this enables stimulation of RAD51 D-loop reactions on RPA-coated ssDNA substrates. Unexpectedly, we find that SWSAP1-SWS1 stimulates RPA diffusion on ssDNA. We find that CRISPR/Cas9 *SWSAP1* and *SWS1* knockout cells are sensitive to the PARP inhibitor, Olaparib, and APE1 inhibition. Importantly, SWSAP1 is mutated in breast, uterine/endometrial and prostate cancers (TGCA, cBioportal) and mutations in SWSAP1 inhibit its binding to SWS1 by yeast-two-hybrid. This study provides novel mechanistic insights into how the RAD51 paralog complex SWSAP1-SWS1 modulates RAD51 and RPA activities and has implications in cancers containing genetic variants in this pathway.

## Results

### Human Shu complex components, SWSAP1 and SWS1, form a heterodimeric protein complex and bind RAD51

It has been presumed that SWSAP1 and SWS1 form a complex based upon their protein-protein interactions by co-immunoprecipitation and mass-spectrometry. Therefore, we sought to determine if untagged recombinant SWSAP1-SWS1 form a complex under physiological conditions. SWSAP1-SWS1 were purified from SF9 insect cells co-infected with baculovirus (**Figure 1A**). We find that recombinant SWSAP1-SWS1 form a heterodimeric protein complex in solution by size exclusion chromatography, where SWSAP1 (27 kDa) and SWS1 (15 kDa) coelute in the same fractions (**Figure 1B**; B12, B11, B10). The protein identity was confirmed using western blot analysis and mass-spectrometry peptide analysis. By generating a standard curve using known low and high size exclusion protein standards, we determined the SWSAP1-SWS1 partition coefficient (K_av_) of 0.53 +/- 0.00 and calculated a molecular weight of 31.58 +/-0.77 kDa (**Figure 1C**). It should also be noted that like the canonical RAD51 paralogs, SWSAP1, must be co-purified with its binding partner, SWS1^2,14,15^. Together, these results demonstrate that SWSAP1-SWS1 form a heterodimeric protein complex in solution under near physiological pH (7.5), salt (150 mM), and ATP (1 mM) concentrations.

**Figure 1:**
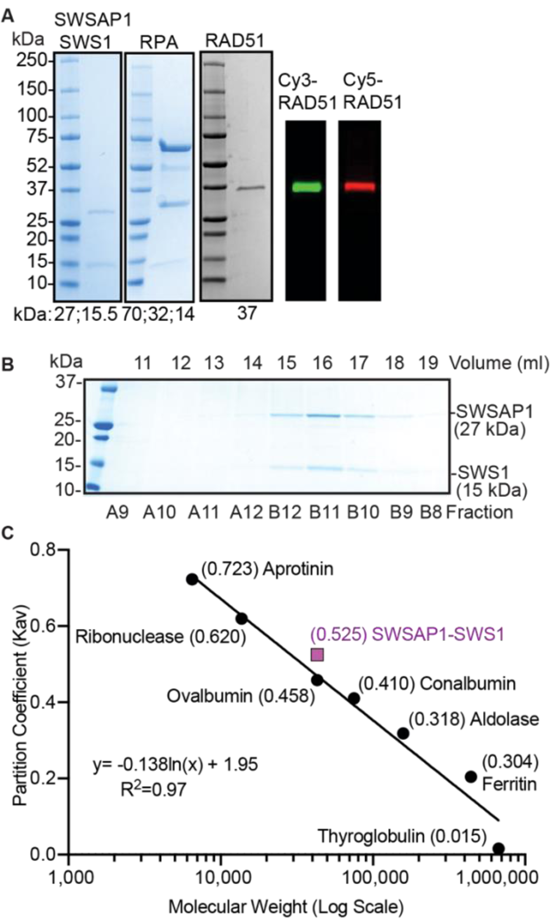
Recombinant SWSAP1-SWS1 form a heterodimeric protein complex that binds RAD51. A) Recombinant SWSAP1-SWS1 (25, 15.5 kDa, respectively), RPA (70, 32, 14 kDa), RAD51 (37 kDa), Cy3-RAD51 (37 kDa), and Cy5-RAD51 (37 kDa) were run on SDS-PAGE gel and Coomassie stained or visualized by typhoon imaging (Cy3-RAD51 and Cy5-RAD51). B) SWSAP1-SWS1 were analyzed by size exclusion and the eluted fractions were analyzed by SDS-Page gel and Coomassie staining. Experiments were performed in triplicate. C) Quantitative analysis was performed to determine the size of SWSAP1-SWS1 (purple text) using known molecular standards and the partition coefficient was graphed vs the log of the molecular weight. The molecular standards used include Aprotinin (6.5 kDa), Ribonuclease A (13.7 kDa), Ovalbumin (43 kDa), Conalbumin (75 kDa), Aldolase (158 kDa), Ferritin (440 kDa), and Thyroglobulin (669 kDa) run on a Superdex200 (10/300 GL) column. Partition coefficients (Kav) of low and high molecular weight protein standards are plotted vs the Log of the MW. Experiments were performed in triplicate.

Next, we investigated the interaction of recombinant RAD51 with the SWSAP1-SWS1 heterodimer. To uncover the function of SWSAP1-SWS1 in RAD51-dependent strand exchange, we purified RAD51, the trimeric replication protein A complex (RPA; consisting of 70, 32, and 14 kDa subunits), and Cy3/Cy5-RAD51 (**Figure 1A**). To determine if SWSAP1-SWS1 binds to RAD51 in the absence of DNA, we incubated SWSAP1-SWS1 (4 and 6 μM) with RAD51 (7.5 μM) and ran these complexes on blue native PAGE gels followed by Coomassie staining (**Supplementary** Figure 1). We found under Mg^2+^ and ATP conditions, where RAD51 forms irregularly-sized filaments, SWSAP1-SWS1 and RAD51 display diminished migration, which is indicative of complex formation (**Supplementary** Figure 1A)^16^. Mass spectrometry was used to confirm that both proteins were present in these higher migrating species (highlighted with white star). These findings are consistent with previous studies demonstrating a direct interaction between RAD51 and SWSAP1 and/or SWS1^8,17^. Together these results support the notion that SWSAP1-SWS1 and RAD51 form protein complexes in solution.

### SWSAP1-SWS1 maintains RAD51-ssDNA integrity and forms interspersed filaments

RAD51 function during homologous recombination and at stalled replication forks relies upon the ability of RAD51 to form a nucleoprotein filament on ssDNA. In the presence of ATP, Mg^2+^ and Ca^2+^, RAD51 forms a right-handed nucleoprotein filament, also known as a pre-synaptic filament, that is stabilized under conditions where ATP hydrolysis is inhibited by Ca^2+ 16,18–21^. Importantly, as RAD51 nucleates on ssDNA it stretches the ssDNA 1.6-fold beyond the dsDNA B-form^11,19,22–28^. This conformation is important for the homology search and strand exchange reactions and varies in size from 300 nucleotides to ∼2.6 kb. To determine if SWSAP1-SWS1 further modulates RAD51 binding and extension on ssDNA substrates, we utilized a Förster Resonance Energy Transfer (FRET)-based ssDNA binding assay^29–31^. This assay utilizes a dT60 substrate (10 nM final) containing an internal Cy3 fluorescent dye (FRET donor) that is separated 25 nucleotides from a Cy5 fluorescent dye (FRET acceptor)^32^. The ssDNA alone adopts a conformation where the two dyes are in close proximity resulting in a high FRET signal. As RAD51 is titrated in the reaction, we observe a decrease in the FRET due to RAD51 binding and stretching the ssDNA, which exhibits an expected stoichiometric binding isotherm as previously reported (**Figure 2A**;^29–31,33^). In agreement with previous findings, RAD51 binds ssDNA with a stoichiometry of three nucleotides per RAD51 monomer (**Figure 2A**; black circles)^29,33^. The stoichiometry was calculated by multiplying the concentration of dT60 (10 nM) by the total number of nucleotides (60) resulting in 600 nM nucleotides. Then, the 600 nM nucleotides was divided by the saturation point (200 nM RAD51) resulting in a stoichiometry of 3 nucleotides bound per RAD51 molecule^29,33^. As a control, we titrated SWSAP1-SWS1 with dT60 and do not observe perturbation of the dT60 substrate under these reaction conditions, suggesting that SWSAP1-SWS1 does not bind ssDNA with high affinity (**Figure 2B**; pink circles).

**Figure 2:**
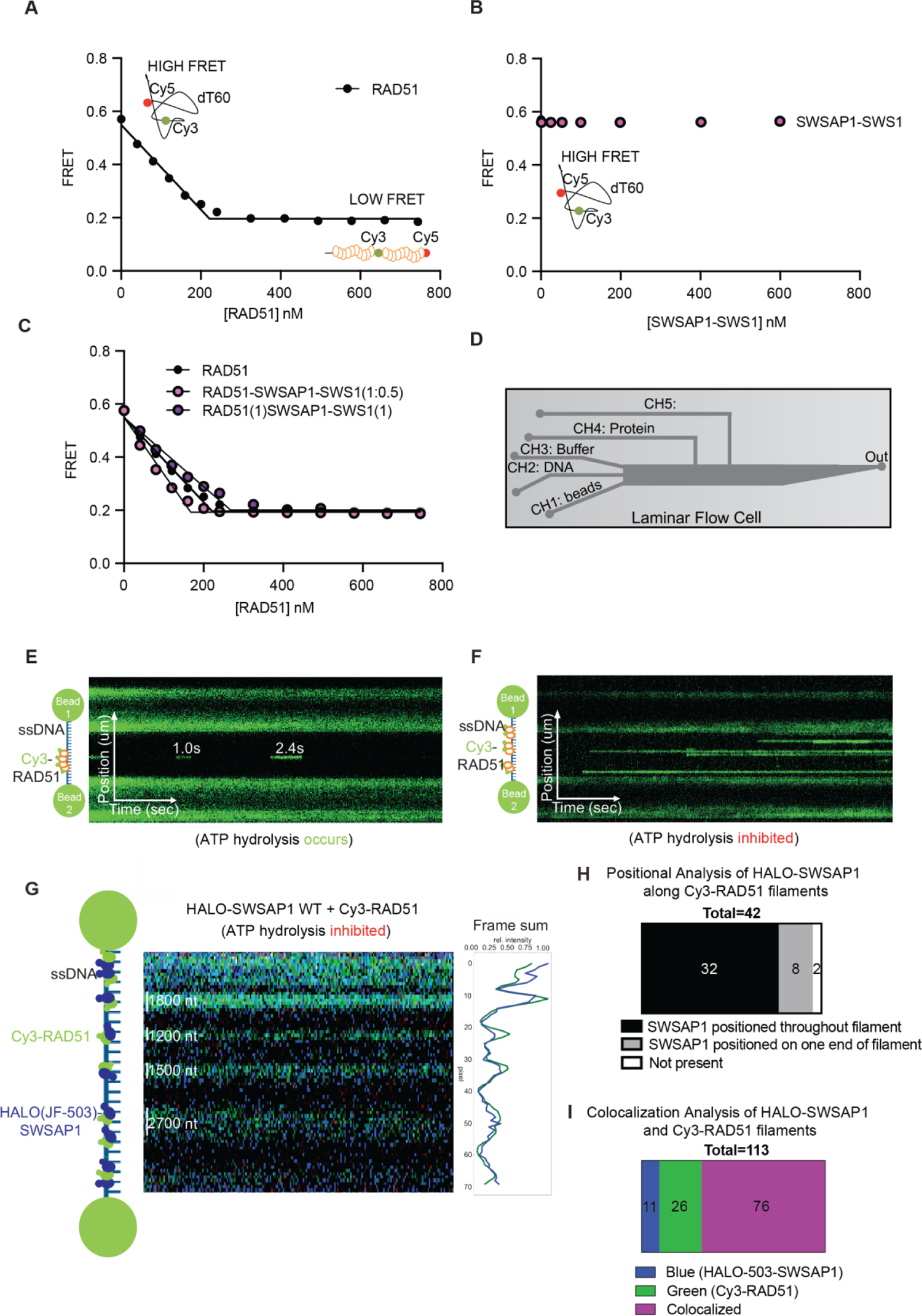
SWSAP1-SWS1 maintains RAD51-ssDNA binding stoichiometry and filament assembly via decorating the outside of the RAD51 filament. A) FRET-based ssDNA binding assays were performed with a dT60 substrate with a Cy3 dye (FRET donor) and a Cy5 dye (FRET acceptor) separated by 30 nucleotides and increasing concentrations of RAD51 (black circles). Experiments were performed in triplicate. B) FRET-based ssDNA binding assays were performed with a dT60 substrate with a Cy3 dye (FRET donor) and a Cy5 dye (FRET acceptor) separated by 30 nucleotides and increasing concentrations of SWSAP1-SWS1 (pink circles). Experiments were performed in triplicate. C) FRET-based ssDNA binding assays with Cy3-dT60-Cy5 were performed with increasing concentrations of RAD51 alone or with SWSAP1-SWS1 at two different molar ratios (1:0.5 molar ratio in pink and 1:1 molar ratio in purple). Experiments were performed in triplicate. D) Schematic of experimental setup for single-molecule studies of Cy3-RAD51 and Cy3-RAD51-HALO-SWSAP1-SWS1 experiments. In channel one lasers trap the streptavidin beads (CH1: beads), then are moved to the second laminar flow channel which bind to biotin-labeled dsDNA (CH2: DNA), then ssDNA is formed in channel three (CH3: Buffer) and moved to channel four containing Cy3-RAD51 or Cy3-RAD51-HALO-SWSAP1-SWS1 nuclear extract proteins (CH4: Protein). E) Representative kymograph showing Cy3-labeled RAD51 on lambda ssDNA under magnesium conditions. Cy3-RAD51 position was monitored over time. In the presence of Mg2+, Cy3-RAD51 ATP hydrolysis occurs resulting in dwell times of ∼1.5 seconds. F) Representative kymograph showing Cy3-labeled RAD51 on lambda ssDNA under calcium conditions. The Cy3-labeled RAD51 and its position was monitored over time. In the presence of Ca2+, Cy3-RAD51 ATP hydrolysis is inhibited resulting in increased dwell times and photobleaching time of ∼11 minutes. G) Representative kymograph showing Cy3-labeled RAD51 on lambda ssDNA complexed with HaloTag-JF503-SWSAP1 from nuclear extracts under calcium conditions. Positions of both RAD51 and SWSAP1 were monitored over time. Positional analysis and summation of fluorescent intensity was graphed relative to pixels in kymograph. The length of the filament is indicated on the left of the image. H) Each scan in the kymograph was summed and quantification of positional localization of SWSAP1 relative to the RAD51 filament [throughout the RAD51 filament (32 total; black bar), on the end of the RAD51 filament (8 total; grey bar), or SWSAP1 not present (2 total; white bar)] n=42. I) The Cy3-RAD51 filaments were traced and quantified for colocalization with HaloTag-JF503-SWSAP1. Of the 113 filaments events observed, RAD51 and SWSAP1 colocalized in 76 events (purple bar) whereas 26 events had RAD51 (green bar) and 11 events had SWSAP1 alone (blue bar).

This result is not unexpected as a related RAD51 paralog containing complex, BCDX2, does not bind ssDNA under physiological salt conditions^2^. To determine if SWSAP1-SWS1 has a function in dismantling RAD51 filaments, we performed binding assays where two different molar ratios of SWSAP1-SWS1 to RAD51 were preincubated together and then FRET was assessed following dT60 addition. We do not observe significant perturbation of the RAD51-ssDNA binding in the presence of SWSAP1-SWS1 (**Figure 2C**). These results contrast with those previously reported for SWSAP1 alone where FIGNL1 and SWSAP1 dismantle RAD51 filaments^8^. Here, we demonstrate that binding of SWSAP1-SWS1 to RAD51 maintains the integrity of the RAD51-ssDNA interaction.

To determine where SWSAP1-SWS1 are positionally located within RAD51 filaments, we utilized LUMICKS C-trap which combines single molecule confocal fluorescence microscopy with optical tweezers. We used the C-trap since it allows large RAD51 filaments to be visualized (up to 2,700 nucleotides in size). This approach more closely recapitulates what occurs after DNA end resection and what is utilized during homologous recombination. We first examined RAD51 alone by generating a Cy3-N-terminally labeled RAD51. We validated that these N-terminally tagged RAD51 proteins form filaments comparable to untagged RAD51 by blue native PAGE gel (**Supplementary** Figure 1B). As expected, Cy3-RAD51 proteins form nucleoprotein filaments of irregular shapes and sizes under magnesium conditions (**Supplementary** Figure 1B). Next, we used a microfluidic flow-cell (LUMICKS) which has five distinct flow channels separated by laminar flow in which the two optical traps are moved through (**Figure 2D**). Due to laminar flow, channels 1-3 remain separated even though there is no physical barrier between them. First, streptavidin-coated 4.38 micron polystyrene beads were captured and biotinylated lambda dsDNA (20,452 nt) (LUMICKS) was captured between the beads. Subsequently, ssDNA was generated as previously described^18,34,35^ and dipped into reservoir 4 containing Cy3-RAD51 alone (80 nM final). The sample was then moved into channel 3 containing imaging buffer and protein-ssDNA interactions were recorded by kymograph over time without flow. To further validate that our Cy3-RAD51 protein is functional, we examined Cy3-RAD51 binding to ssDNA under conditions where ATP hydrolysis is regulated by Mg^2+^ or Ca^2+^. As previously shown, we observe shorter RAD51 filament dwell times under conditions which allow for ATP hydrolysis (Mg^2+)^ with a duration of “ON states” of ∼1.5 seconds [**Figure 2E**; ^21,36^]. When ATP hydrolysis is inhibited (Ca^2+^), we observe enhanced RAD51 filament dwell times of 11 minutes [**Figure 2F**; ^21^]. Together, these results demonstrate that Cy3-labeled RAD51 protein is functional.

To determine where SWSAP1-SWS1 binding occurs along the RAD51 filament, we performed C-trap assays with HaloTag-JF503-SWSAP1 protein (2-15 nM) from RPE-1 nuclear extracts and Cy3-RAD51 (80 nM final) (**Figure 2G**)^34^. Note that HaloTag-SWSAP1 and HA-SWS1 were co-transfected into *sgSWSAP1* RPE-1 knockout cells so that only the HaloTag-SWSAP1 was expressed and JF503 was subsequently added to generate fluorescent SWSAP1 protein. We verified expression of HaloTag-SWSAP1 by exciting the SDS-PAGE gel at 488 nm on the Typhoon (**Supplementary** Figure 1C**, 1D**). We used this approach due technical difficulties associated with making fluorescently labeled complexes *in vitro* and so that post-translational modifications and other important co-factors would be present in the extract^34^. We examined the position of both fluorescent proteins on ssDNA over time. Surprisingly, positional analysis revealed that SWSAP1 is primarily decorated throughout the Cy3-RAD51 filaments (32 out of 42; **Figure 2H**). In support of the positional analysis, HaloTag-JF503-SWSAP1 colocalized with the majority of RAD51 filaments analyzed 67% of the time (76 out of 113; **Figure 2I**). Together, our results show that SWSAP1-SWS1 binds RAD51, where it is located throughout the filament. Note that using this approach, SWSAP1-SWS1 could be incorporated within the RAD51 filament itself or decorated along the outside of the RAD51 filament through its interaction with RAD51.

### SWSAP1-SWS1 stimulates RAD51-mediated D-loop formation on RPA-coated ssDNA

To determine if SWSAP1-SWS1 interaction with RAD51 is important for RAD51’s function in homology search and strand exchange, we examined whether SWSAP1-SWS1 stimulates RAD51-mediated displacement-loop (D-loop) formation. In this assay, a 90 nt ssDNA FAM-labeled substrate is used that is complementary to a sequence in the pBlueScript KG dsDNA substrate. First, the FAM-90 mer ssDNA is coated with RPA and subsequently, RAD51 and/or SWSAP1-SWS1 is added at increasing concentrations (**Figure 3A**). D-loop formation occurs when the pBlueScript KG dsDNA plasmid, which contain homology to the ssDNA, is added to the reaction. The reaction products are separated on an agarose gel and the D-loop product is quantified relative to RAD51-RPA alone (**Figure 3B, 3C**). We examined whether SWSAP1-SWS1 stimulates RAD51 D-loop formation on RPA-coated ssDNA. We observe that SWSAP1-SWS1 stimulates RAD51-mediated D-loop formation (**Figure 3B, 3C**). We observe an increase in D-loops formed from SWSAP1-SWS1 (1.25 nM to 250 nM), which resulted in up to ∼4-fold stimulation of D-loops (**Figure 3C**). We also examined whether the non-cleaved SWSAP1 and GST-tagged SWS1 complexes stimulate D-loop formation and find up to 2-3-fold stimulation depending on the concentration of GST-SWS1-SWSAP1 (**Supplementary** Figure 2A**, 2B**). Therefore, the GST tag does partially attenuate GST-SWS1 activity compared to the untagged protein. Importantly, GST-SWS1-SWSAP1 and GST-SWS1-SWSAP1-RPA complexes do not promote D-loop formation alone (**Supplementary** Figure 2C**, 2D**). These results demonstrate that both constructs with and without recombinant tags on SWSAP-SWS1 can stimulate RAD51 strand exchange and D-loop formation when challenged with RPA-coated ssDNA.

**Figure 3:**
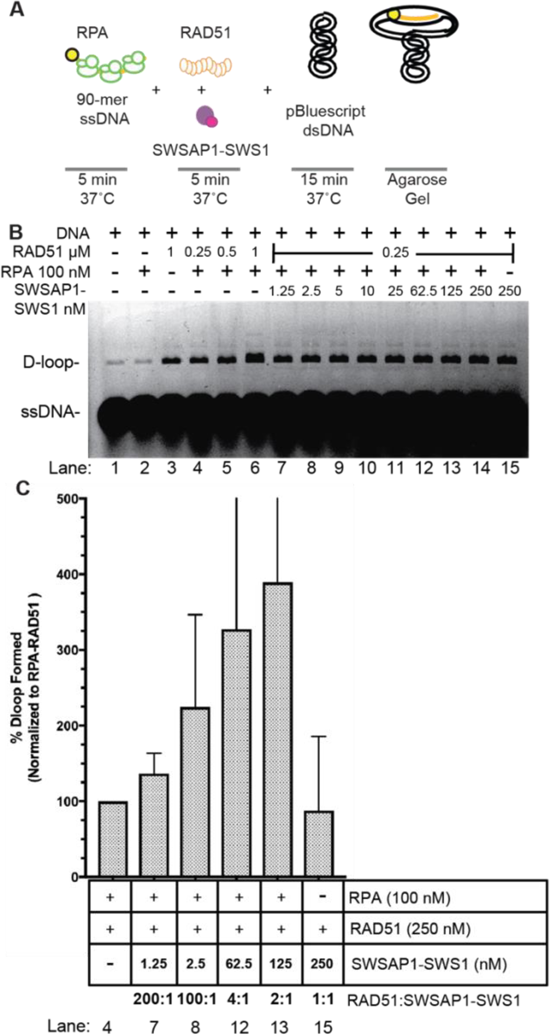
SWSAP1-SWS1 stimulates RAD51 D-loop activity on RPA-coated ssDNA. A) Schematic depicting D-loop assay. FITC-90mer ssDNA (yellow DNA with circle) was coated with RPA (green circles) for 5 min at 37°C. Subsequently, RAD51 (orange circles), SWSAP1-SWS1 (pink and purple circles), or RAD51 with SWSAP1-SWS1 was added to the reaction for 5 min at 37°C and then incubated with the pBluscript dsDNA template (supercoiled DNA) for 15 min at 37°C. D-loops were deproteinated, run on an agarose gel, and imaged for FITC signal. B) FITC-90mer ssDNA coated with RPA (100 nM) was challenged with either RAD51 (0.25, 0.5, 1 μM) or RAD51 (0.25 μM) and increasing concentrations of SWSAP1-SWS1 as indicated. Then the reaction was begun by adding pBluescript dsDNA. D-loops were deproteinated, run on an agarose gel, and imaged for FITC signal. DNA alone, DNA + RPA, DNA + RAD51, DNA + RAD51 + RPA are controls (Lanes 1-6). C) Quantification of D-loop reaction products with recombinant SWSAP1-SWS1. The reactions utilized 100 nM RPA, 250 nM RAD51, and SWSAP1-SWS1 (1.25, 2.5, 62.5, and 125 nM) proteins and the values were normalized to RPA 100 nM and RAD51 250 nM. Experiments was performed five times and standard deviations are shown.

### SWSAP1-SWS1 binds RPA and enhances RPA diffusion on ssDNA

It is puzzling that SWSAP1-SWS1 does not modulate RAD51-ssDNA filament stability but can stimulate RAD51 activities. Therefore, we hypothesized that SWSAP1-SWS1 may stimulate RAD51 through a functional interaction with RPA. To determine if SWSAP1-SWS1 binds RPA in the absence of DNA, we performed mass photometry experiments. We observe SWSAP1-SWS1 (1 μM) control at 48 kDa and RPA (25 nM) at 110 kDa (**Figure 4A**). When SWSAP1-SWS1 and RPA are mixed together, we observe disappearance of the SWSAP1-SWS1 heterodimer and appearance several larger complexes observed at molecular weights of 127 and 229 kDa (**Figure 4A**). Note that the different estimated molecular weights for SWSAP1-SWS1 when using mass photometry and size exclusion. Unfortunately, the mass photometry technique is limited in quantifying low molecular weight complexes and the limit of the detector is ∼56kDa (beta-amylase) which explains the higher calculated molecular weight observed with mass photometry. These results show that SWSAP1-SWS1 and RPA physically interact in solution in the absence of DNA. To determine if SWSAP1-SWS1 facilitates RPA dissociation from ssDNA, we performed FRET-based ssDNA binding assays using a dT60 substrate (10 nM substrate; **Figure 4B)** as previously described^37^. As RPA has a higher affinity to ssDNA compared to RAD51 (picomolar *K_d_*<10^-10^ M vs nanomolar for RAD51), we hypothesized that SWSAP1-SWS1 may remodel or redistribute RPA on ssDNA to enable RAD51 function. We observe stoichiometric binding with a plateau at 40 nM RPA suggesting at least four RPA molecules are engaging with ssDNA under these reaction conditions (**Figure 4B**)^29,38^. For these experiments 150 mM salt was used to reflect the most physiological concentration that occurs *in vivo*. This salt concentration results in microscopic and macroscopic changes in the domains of RPA that interact with the ssDNA and this can change occupancy as described by biochemical and single-molecule analysis^39^. We challenged the RPA-ssDNA binding interaction (40 nM RPA-10nM dT60) by titrating SWSAP1-SWS1 at increasing concentrations. We observed an increase in the FRET signal upon SWSAP1-SWS1 addition (**Figure 4C**). Interestingly, we do not observe a full change in FRET back to ∼0.58. These data suggest that SWSAP1 may be promoting partial RPA dissociation or remodeling the RPA-ssDNA interaction. The three RPA subunits are comprised of six OB-fold domains connected by flexible linkers, there are numerous ways one could speculate RPA kinetics are modulated^40^. Together, these results show that SWSAP1-SWS1 binds to RPA and remodels the RPA-ssDNA interaction and explains the mechanism of how SWSAP1-SWS1 stimulates RAD51-mediated D-loop formation.

**Figure 4:**
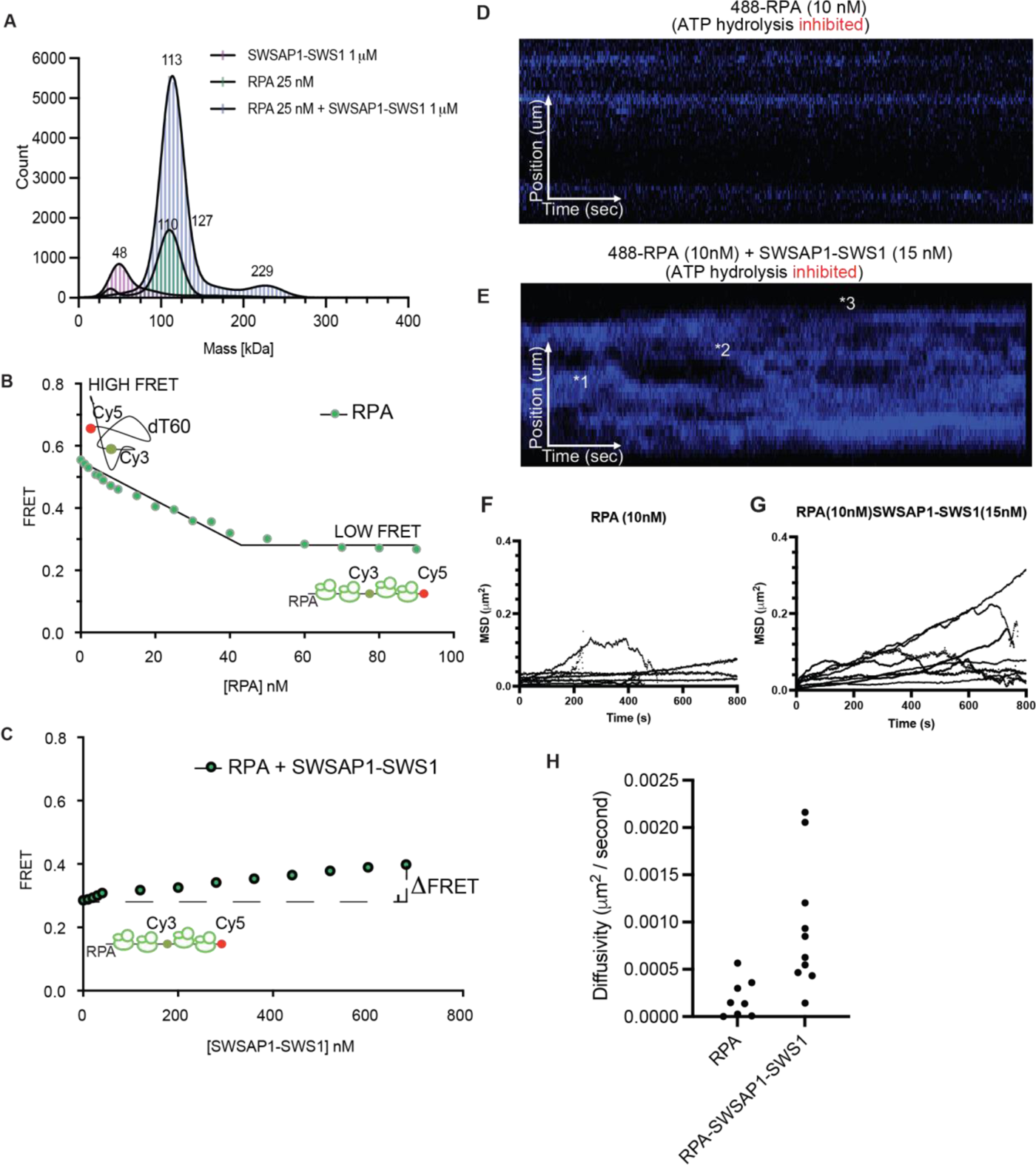
SWSAP1-SWS1 binds RPA and enhances RPA diffusion on ssDNA. A) Mass photometry analysis show that SWSAP1-SWS1(1 μM) (48 kDa) binds to RPA (25nM) (110 kDa) and forms higher order complexes. Experiments were performed in triplicate. Note that the different estimated molecular weights for SWSAP1-SWS1 when using mass photometry and size exclusion. Unfortunately, the mass photometry technique is limited in quantifying low molecular weight complexes and the limit of the detector is ∼56kDa (beta-amylase) which explains the higher calculated molecular weight observed with mass photometry. B) FRET-based ssDNA binding assay where substrate (10 nM dT60) titrated with increasing concentrations of RPA (green) show stoichiometric binding isotherm with 4 RPA molecules. Experiments were performed in triplicate. C) FRET-based ssDNA binding assays were saturated with RPA-ssDNA complexes (40 nM RPA, 10 nM dT60) and challenged with increasing concentrations of SWSAP1-SWS1 protein. Experiments were performed in triplicate. D) Representative kymograph showing AlexaFluor488-RPA protein on lambda ssDNA over time. The lambda DNA was moved into a channel containing the AlexaFluor488-RPA protein (10 nM) (blue) and imaged in buffer. Positions of AlexaFluor488-RPA protein was monitored over time. Positional analysis, mean squared displacement, and diffusivity was calculated. E) Representative kymograph showing AlexaFluor488-RPA protein incubated with SWSAP1-SWS1 on lambda ssDNA over time. The lambda DNA was moved into a channel containing the AlexaFluor488-RPA protein (10 nM) (blue), into buffer, and then into a channel containing unlabeled SWSAP1-SWS1 (15nM) for imaging. Positional analysis, mean squared displacement, and diffusivity was calculated. *1, *2, and *3 are examples of dynamic molecules which have undergone large displacements (5 kb or >) at long time scales. F) MSD analysis of the trajectories was graphed vs time for AlexaFluor488-RPA protein alone (n=21). G) MSD analysis of the trajectories was graphed vs time for AlexaFluor488-RPA (n=21) protein or AlexaFluor488-RPA protein and SWSAP1-SWS1 (n=29). H) Diffusivity in microns^2^ per second were graphed for each MSD in either AlexaFluor488-RPA (n=21) or for AlexaFluor488-RPA protein and SWSAP1-SWS1 (n=29).

To determine if SWSAP1-SWS1 is changing RPA dynamics on ssDNA, we again utilized a single molecule confocal fluorescence microscopy combined with optical tweezers setup to visualize AlexaFluor488-RPA on ssDNA under tension of 10 or 25 pN^41,42^. Briefly, ssDNA was generated, the beads and DNA complex were moved to a channel 4 containing 10 nM AlexaFluor488-RPA (10nM) for 10 seconds, and the complex was moved back into buffer for kymograph imaging. As expected, AlexaFluor488-RPA protein binds to lambda ssDNA (**Figure 4D**) and mean square displacement was calculated and graphed vs time (**Figure 4F**). On long timescales, RPA alone diffuses very little at forces of 10 or 25 pN as previously shown (**Figure 4F)**^43^. To determine if SWSAP1-SWS1 binding enhances dynamics, which would be observed as an increased mean square displacement and calculated RPA diffusivity, we again created AlexaFluor488-RPA complexes on ssDNA in channel 4, and moved the protein-DNA complexes into channel 5 containing 15 nM unlabeled SWSAP1-SWS1 protein without flow thus allowing for visualization of RPA diffusion (**Figure 4E**).

Surprisingly, RPA molecules began diffusing on ssDNA in the presence of SWSAP1-SWS1. When the mean square displacement of RPA was graphed, we observe many dynamic molecules which have undergone large displacements (5 kb or >) at long time scales (**Figure 4G**; Star 1,2,3* **Figure 4E**). Each pixel represents 300 nucleotides of ssDNA. Interestingly, we also observe events in which new RPA filaments appear, suggesting that SWSAP1-SWS1 not only facilitates RPA diffusion, but may also promote hopping. To quantitatively describe this phenomenon, we calculated and compared the diffusivity of AlexaFluor488-RPA and AlexaFluor488-RPA when incubated with SWSAP1-SWS1 (**Figure 4H**). Again, we observe a large enhancement of RPA diffusion in the presence of SWSAP1-SWS1. Together, these data suggest that SWSAP1-SWS1 binding to RPA may cause some change that enhances the kinetic macroscopic and/or microscopic on and off rates of RPA on ssDNA.

### SWSAP1 mutations found in breast, uterine, endometrial, and prostate cancers disrupt binding to SWS1 and *sgSWSAP1 sgSWS1* cells are sensitive to PARPi, Olaparib

Disruption of *SWSAP1* and *SWS1* in humans is associated with fertility defects and mutations in *SWSAP1* has been identified in cancer databases including cBioPortal and The Cancer Genome Atlas (TCGA)^44–46^. Similarly, mutations in the canonical *RAD51* paralogs, including *RAD51C* and *RAD51D*, have been identified in hereditary cancers such as breast, ovarian, and prostate^22,47,48^. Intriguingly, many of these cancer variants result in loss of protein interactions between the RAD51 paralogs and their complex members, which is predictive of loss of homologous recombination function^22,47,48^. Using cBioPortal and The Cancer Genome Atlas (TCGA) databases, we identified 16 cancer variants in *SWSAP1* from breast, uterine, endometrial, and prostate cancers (**Figure 5A**). We analyzed these SWSAP1 variants for altered protein interaction with SWS1 by yeast-two-hybrid and found that 11 of the 16 variants had reduced interaction with SWS1 (**Figure 5B**). Unfortunately, we were unable to verify expression of the SWSAP1 variants by western blot with commercial antibodies, antibodies that we generated, or with the BD antibody. Therefore, we cannot rule out that the variants impact protein expression. We used alpha-fold to generate individual models of SWSAP1 and SWS1 proteins and fed these PDB generated models into the HDOCK server to obtain a theoretical protein-protein interaction model (**Figure 5C**). The HDOCK server creates protein-protein interaction models based on a hybrid algorithm of template-based modeling and *ab initio* free docking. The best model we obtained predicted five of the variants to be surface exposed and near a potential interface with SWS1 (**Figure 5C**, residues in red;^49,50^).

**Figure 5:**
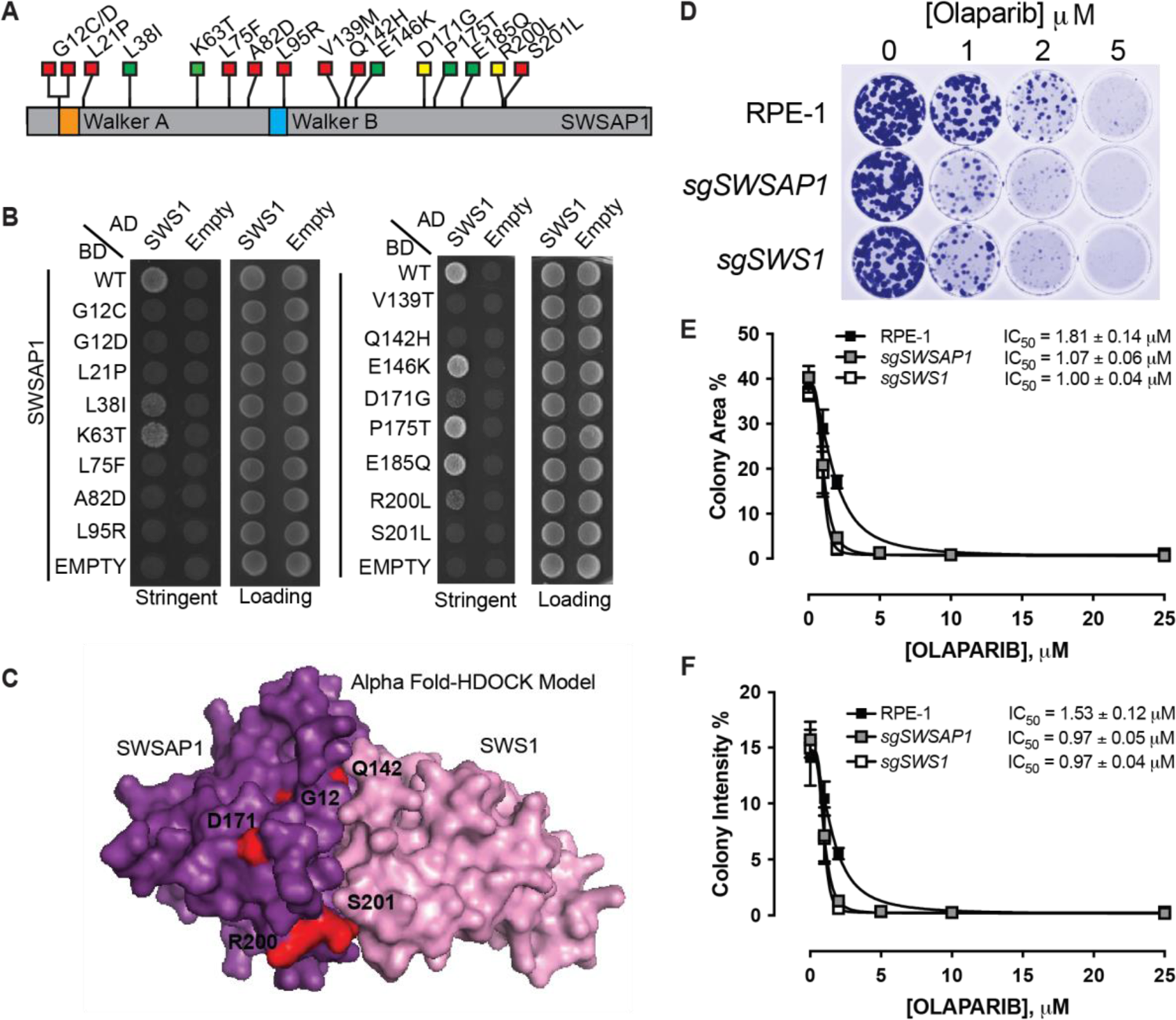
*sgSWSAP1* and *sgSWS1* cells are sensitive to Olaparib inhibition, while breast, uterine/endometrial and prostate cancer mutations inhibit protein-protein interactions. A) Schematic of the SWSAP1 protein, which is 229 amino acids, highlighting variants identified in breast, uterine/endometrial, and prostate cancers from TCGA and cBioPortal. The Walker A and Walker B motifs are indicated in orange and blue, respectively. Summary of the yeast-2-hybrid results in (B) are highlighted with a red box (indicating a deficient yeast-2-hybrid interaction, 0-33%), yellow box (indicating a partial yeast-2-hybrid interaction, 34-67%), and a green box (indicating a proficient yeast-2-hybrid interactions, 68-100%) between SWSAP1 with SWS1. B) Yeast-two-hybrid analysis of SWSAP1 variants from breast, uterine/endometrial, and prostate cancers identified in COSMIC and cBioPortal. Wild-type SWSAP1 or the cancer variant were expressed in the GAL4-DNA binding domain vector (BD) and their interaction with SWS1, expressed in the GAL4-DNA activating domain (AD) was assessed by yeast-two-hybrid by assessing growth on selective medium (SC medium minus leucine, tryptophan, and histidine; SC-LTH). Equal cell plating was assessed by plating the yeast on SC medium minus leucine and tryptophan (SC-LT) medium. pGAD and pGBD empty vectors were used as negative controls. Experiments were performed in triplicate and colonies quantified with ImageJ and normalized to the wild-type control. C) AlphaFold models of SWSAP1 (purple) and SWS1 (pink) structures were fed into the HDOCK server to generate a combined model of SWSAP1-SWS1. The surface exposed cancer variants are highlighted in red. D) Clonogenic survival assays of RPE-1, *sgSWSAP1*, and *sgSWS1* RPE-1 cells exposed to the indicated dose of Olaparib for approximately 14 days. The cells were fixed and stained with crystal violet before being photographed. E) Quantification of (D) where cellular survival area of RPE-1, *sgSWSAP1*, and *sgSWS1* was normalized to untreated control and the colony area quantitated and the IC_50_ was calculated in Graphpad PRISM. Significance was determined by two-way ANOVA from three experiments (SWS1-C1 p=<0.0001, SWS1-C2 p=<0.0001, SWSAP1-C1 p=<0.0001, SWSAP1-C2 p=<0.0001). F) Quantification of (D) where colony intensity of RPE-1, *sgSWSAP1*, and *sgSWS1* was normalized to untreated control and the colony intensity quantitated and the IC_50_ was calculated in Graphpad PRISM. Note that two independent clones were analyzed for each knockout. Significance was determined by two-way ANOVA from three experiments (SWS1-C1 p=<0.0001, SWS1-C2 p=<0.0001, SWSAP1-C1 p=<0.0001, SWSAP1-C2 p=<0.0001).

Homologous recombination-deficient tumors are sensitive to the PARP inhibitor, Olaparib^51^. Importantly, PARP inhibitors have been found to covalently link PARP1 to abasic sites^51,52^. Therefore, we asked whether loss of *SWSAP1* or *SWS1* would result in Olaparib sensitivity. We performed clonogenic survival assays in two independent *SWSAP1* and *SWS1* knockout RPE-1 cells treated with increasing concentrations of Olaparib. We find that *sgSWS1* and *sgSWSAP1* RPE-1 cells are modestly but significantly sensitive to Olaparib (**Figure 5D**). When we quantitated the colony size, we observe approximately a two-fold decrease in colony area (and colony intensity) in cells compared to the parental RPE-1 cell line (**Figure 5E, 5F;** SWS1-C1 p=<0.0001, SWS1-C2 p=<0.0001, SWSAP1-C1 p=<0.0001, SWSAP1-C2 p=<0.0001 for both graphs). These results suggest that loss the human Shu complex, like the other RAD51 mediators, sensitizes cells to PARP inhibition.

## Discussion

The human Shu complex is composed of the RAD51 paralog, SWSAP1, and SWS1. SWSAP1 and SWS1 also interact with SPIDR and PDS5B but the significance of these protein interactions remains unknown^6^. Despite the discovery of the human Shu complex over 15 years ago, the biochemical function of the Shu complex during homologous recombination has remained elusive. We find that SWSAP1-SWS1 binds to and decorates RAD51 filaments on ssDNA where it promotes RAD51 filament assembly. Surprisingly, we find that SWSAP1-SWS1 binds RPA and modulates RPA diffusion on ssDNA, thus enabling RAD51-mediated strand exchange activities. These results uncover the underlying mechanism of how the Shu complex functions to promote RAD51 and RPA activities, which is unique from the other RAD51 paralogs, which cap the filament ends^53^. Our findings provide a rationale as to why so many RAD51 paralog containing complexes exist and suggest that the Shu complex function is distinct from other RAD51 paralog complexes.

Here we uncovered two mechanisms by which the SWSAP1-SWS1 functions to promote homologous recombination. In the first mechanism, we observe SWSAP1-SWS1 binds directly to RAD51 independently of ATP hydrolysis. These results are supported by previous findings examining SWSAP1 and RAD51 interactions^5,8,17^. Using FRET-based biophysical ssDNA binding assays and blue native-PAGE, we show that SWSAP1-SWS1 binds to RAD51. Although SWSAP1-SWS1 binding maintains RAD51 stoichiometry and filament formation on DNA, and SWSAP1-SWS1 is observed throughout RAD51 filaments in the C-trap analysis. RAD51-SWSAP1-SWS1 filaments may be functioning on ssDNA during gap repair to enable multiple nucleation sites for repair. Our previous work showed that *sgSWSAP1* and *sgSWS1* knockout cells have replication restart defects^6^. Therefore, our observation of mixed filaments suggests that SWSAP1-SWS1 may increase the number of RAD51 nucleation sites possible for RAD51-templated repair during ssDNA gaps repair at stalled replication forks. It is also plausible that binding of SWSAP1-SWS1 may change the conformation of the RAD51 protein on ssDNA resulting in a more proficient conformation for strand exchange. This would result in a more “open” or stretched state^23,24^. Like other biochemical studies with the canonical RAD51 paralog complexes, BCDX2 and CX3, purification of functional SWSAP1-SWS1 in our laboratory requires buffers containing ATP and magnesium^2^.

Combined together with our single-molecule optical tweezer data, we show that SWSAP1-SWS1 binds RAD51 positionally decorated throughout the RAD51 filament independently of RAD51 enzymatic activity. Our work with the human Shu complex is distinct from the *C. elegan*s RAD-51 paralogs, RFS-1/RIP-1 ^54^. In *C. elegans* RFS-1 and RIP-1 bind to the SWS1 homolog, SWS-1 ^55^. Unlike SWSAP1-SWS1, RFS-1 and RIP-1 preferentially bind to the 5’ end of RAD-51 filaments ^54^. It is plausible that binding to the end of a filament suggests inhibition of RAD51 disassembly, which is divergent from our FRET-based binding assays with SWSAP1-SWS1 and RAD51 which enable RAD51 filament assembly. One important difference between our studies and the RFS-1/RIP-1 work is that ssDNA was used instead of a dsDNA/ssDNA hybrid. Furthermore, a limitation of our single molecule analysis is that other cofactors are present in the nuclear extracts used in the experiment. Future studies using recombinant proteins will enable independent validation of our findings.

A second mechanism by which SWSAP1-SWS1 stimulates RAD51-mediated D-loop activity is through a DNA-independent physical interaction with RPA. We find that SWSAP1-SWS1 enhances RAD51-mediated D-loop formation only on RPA-coated substrates (**Figure 6**). Importantly, SWSAP1-SWS1 with RPA does not enhance D-loop formation. Using biophysical FRET-based equilibrium ssDNA binding assays, we uncovered that SWSAP1-SWS1 is modulating how RPA interacts with ssDNA. In these studies, we surprisingly observed that the FRET signal does not return to a high FRET state upon SWSAP1-SWS1 addition suggesting SWSAP1-SWS1 may be only interacting with one of the RPA subunits. These findings are supported by the C-trap tweezer experiments examining RPA dynamics in real time in the presence of recombinant SWSAP1-SWS1. Intriguingly, when RPA-ssDNA tethers are incubated in the presence of unlabeled SWSAP1-SWS1 without flow we observed both enhanced diffusive motion of RPA along the DNA. These findings suggest that SWSAP1-SWS1 is modulating RPA dynamics as multiple RPA binding and unbinding cycles can be observed within one kymograph. While it is important to understand how SWSAP1-SWS1 binds RPA, this is complicated by RPA being a heterotrimeric complex with multiple shared ssDNA binding site domains^39,40,56^. Overall, our data suggests that SWSAP1-SWS1 enhances RPA diffusion on ssDNA.

**Figure 6:**
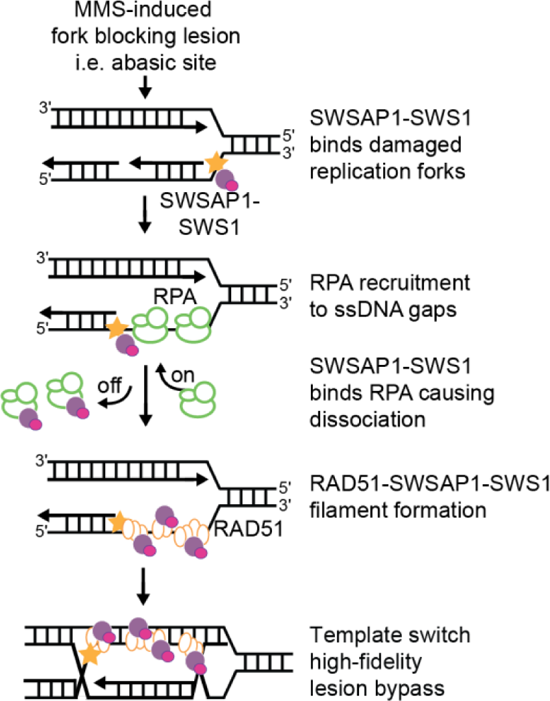
Two mechanisms by which SWSAP1-SWS1 protein stimulates RAD51-mediated high-fidelity repair on RPA coated ssDNA. Schematic of our working model where replicating cells exposed to MMS cause fork blocking lesions, such as abasic sites, within lagging strands of replication forks. The Shu complex binds to MMS-induced DNA damage or processing intermediates, which can stall replication leading to ssDNA gaps behind the fork. RPA-coated ssDNA protects the fork from collapse. SWSAP1-SWS1 interacts with RPA enabling its dissociation from the remodeled fork. RPA dissociation by SWSAP1-SWS1 would then promote RAD51-SWSAP1-SWS1 filament accumulation at these sites to enable template-switching and high-fidelity lesion bypass.

In addition to the function of the RAD51 paralogs in homologous recombination, the RAD51 paralogs also play important functions in the bypass of replicative damage. The BCDX2 and CX3 complexes function in replication fork protection or restart, respectively^1,57^. Similarly, we previously showed that *SWSAP1* and *SWS1* knockout cells also have defective fork restart upon HU treatment^6^. More recently, the BCDX2 complex was linked to fork reversal where the BCDX2 complex stimulates SMARCAL1 and ZRANB3 strand annealing and branch migration activities independently of RAD51^58^. Like the BCDX2 complex, it is also plausible that SWSAP1-SWS1 may have homologous recombination-independent functions during replication stress.

Cancer cells have increased replication stress and cancer therapeutics exploit these genetic vulnerabilities to mediate cell death. Like the other RAD51 mediators, *SWSAP1* and *SWS1* variants have also been identified in tumors and individuals with *SWS1* germline variants have decreased fertility including defects in spermatogenesis and polycystic ovarian syndrome^44–46^. Furthermore, we find that many of the breast, uterine/endometrial, and prostate cancer identified variants exhibit defects in the SWSAP1-SWS1 protein interactions, which is important for complex function. Similar to what has been observed in other homologous recombination deficient tumors, we find that cells with *SWSAP1* and *SWS1* genes knockout are sensitive to Olaparib. This finding is consistent with reduced RAD51 foci formation upon replication stress^6^, which may in part be due to increased ssDNA gaps that form during replication^59^. Loss of the canonical RAD51 paralogs similarly results in PARP inhibitor sensitivity and may predict therapeutic response in pathogenic variants within this protein family^47,60,61^. These results suggest that individuals with pathogenic Shu complex mutations may also be sensitive to homologous recombination-deficient targeted therapies. Furthermore, our preliminary data show that Shu complex deficient cells are modestly sensitive to APE1 inhibition (**Supplementary** Figure 3). Abasic sites can form when the alkylated base, such as three-methyl adenine, is removed by a DNA glycosylase. Subsequently, the APE1 endonuclease processes abasic sites and its inhibition results in abasic site accumulation^62^. These results suggest that SWSAP1-SWS1 are important for cellular survival when abasic sites accumulate. Therefore, the human Shu complex, like the yeast complex, may play a key role in bypassing abasic sites encountered during replication by promoting RAD51 filament formation and template switching (**Figure 6**).

Here we provide evidence that like the other RAD51 paralogs, the Shu complex plays an important and distinct role in RAD51 regulation, and whose disruption may have profound implications for human disease.

## Methods

### Constructs

All constructs used for infection of SF9 Insect Cells (Gibco) were codon optimized and cloned into vector pVL1393 (MTI Bio). The 6XHIS-SWSAP1 and GST-SWS1 constructs were created and optimized for expression in SF9 insect cells (MTI Bio). The pCH1-RAD51 vector was obtained from Dr. Maria Spies (University of Iowa).

### Protein Purification

Human RAD51 and RPA were expressed and purified as described^30,33,37,39^. The AlexaFluor488-RPA protein was provided by Dr. Shixin Liu (Rockefeller University) and purified as described^41,42^. The human SWSAP1 and SWS1 proteins were purified from baculovirus infected SF9 insect cells (MTI Bio). After infection of 2 × 10^7^ SF9 (Gibco) with baculovirus (GST-SWS1 MOI=5 and SWSAP MOI=5) for 48 hrs, cells were centrifuged and pellets stored at −80°C. Pellets were removed from ice, solubilized in 50 mL of lysis buffer (Lysis Buffer: 100 mM Tris pH 8, 300 mM NaCl, 5 mM BME, 10% glycerol, 20% sucrose, 0.01% NP40, 1 mM ATP, 5 mM MgCl2, 1 mM ZnCl2, 2 PhosSTOP easy Pack (ROCHE), Protease Inhibitor cocktail (Sigma), 1 mM PMSF, 30 μg/mL leupeptin) and lysed by sonication. Lysed supernatant was centrifuged for 2 hrs at 45,000 rpm at 4°C. Then the supernatant was incubated with 20 mL of glutathione agarose beads for 1.5 hrs at 4°C. Beads were washed with lysis buffer and low salt buffer (Low Salt Buffer: 40 mM Tris pH 8, 150 mM NaCl, 1 mM BME, 10% glycerol, 0.01% NP40, 1 mM ATP, 5 mM MgCl2). Then protein was eluted with 10 mM reduced glutathione. Elution was then added with 250U biotin labeled thrombin (EMD Millipore) and dialyzed in 3,350 MW dialysis tubing o/n in 1L Thrombin Dialysis Buffer (Thrombin Dialysis Buffer: 40 mM Tris pH 8, 150 mM NaCl, 1 mM BME, 20% glycerol, 1 mM ATP, 5 mM MgCl2) at 4°C. Cleaved GST and uncleaved GST-SWS1 was captured on a GST column by incubating with 20 mL of glutathione agarose beads for 1 hr at 4°C. The elution was collected and added to 10 mL of StrepTactin beads for capture of biotinylated thrombin. Flow through containing cleaved SWSAP1-SWS1 was added to 20 mL of IMAC Sepharose charged with NiS04. The column was washed with 100 mL of 5, 10, and 15 mM Imidazole buffers and eluted with 100 mM Imidazole. Eluted fractions were run on a Superdex 200 size exclusion column in Size Exclusion Buffer (Size Exclusion Buffer: 40 mM Tris pH 8, 300 mM NaCl, 1 mM BME, 10% glycerol, 0.01% NP40, 1 mM ATP, 5 mM MgCl2). The protein eluted as a heterodimer at the expected molecular weight. Protein was concentrated on a Mono Q column and dialyzed o/n into storage buffer (Storage Buffer: 40 mM Tris pH 8, 150 mM NaCl, 1 mM DTT, 40% glycerol, 0.01% NP40, 1 mM ATP, 5 mM MgCl2). Protein was flash frozen and stored at –80°C.

### RAD51 Cy3 and Cy5 N-terminal labeling

RAD51 was N-terminally labeled with either Cy3 or Cy5. Labeling was performed by incubating RAD51 protein that was dialyzed in buffer containing 250 mM NaPi (pH 7.0), 150 mM NaCl, 1 mM DTT, and 10% glycerol with Cy3-Mono-Reactive Dye or Cy5-Mono-Reactive Dye overnight (VWR Catalog# 95017-379). The dye-labeled Cy3-RAD51 and Cy5-RAD51 was further purified as described above to remove any free dye in the protein prep. Labeling efficiency was determined by measuring the absorbance of RAD51 at 280 nm and of Cy5 at 550 nm using their respective (ε_280_ =14,900 M^-1^cm^-1^ for RAD51 and ε_550_ =150,000 M^-1^cm^-1^ for Cy3). Labeling efficiency was determined to be 69% for Cy3-RAD51. Cy5-RAD51 was further purified as described. Labeling efficiency was determined by measuring the absorbance of RAD51 at 280 nm and of Cy5 at 650 nm using their extinction coefficients (ε_280_ =14,900 M^-1^cm^-1^ for RAD51 and ε_650_ =250,000 M^-1^cm^-1^ for Cy5). Labeling efficiency was determined to be 40% for Cy5-RAD51.

### Quantitative Size Exclusion Analysis

Size exclusion analysis was performed using a Superdex 10/300 GL Agarose Column (GE) equilibrated in buffer (20 mM Hepes KOH pH7.5, 150 mM NaCl, 1 mM ATP, 5 mM MgCl_2_, and 1 mM DTT). Molecular weight standards used from the Gel Filtration Calibration Kit HMW (GE/Cytiva) included Dextran, Aldolase, Conalbumin, Ovalbumin, Ferritin and Thyroglobulin were used. Small molecular weight proteins utilized for size calibration included Proteinase K (AMBION), rTEV, Aprotonin (A6279 Sigma), Ribonuclease A, from bovine pancreas (R5125 Sigma). Proteins were run over column at 0.5 mL/min and 1 mL fractions were collected over a geometric column volume (V_c_)of 24.19 with bed height of 30.8mm. Dextran was utilized to calculate the void volume (V_o_) and corresponded to 8.05 mL. The elution volume at max absorbance of the eluted protein (V_e_) was determined and was used to calculate the partition coefficient (K_av_). Kav was calculated by (Kav=(V_e_-V_o_)/(V_c_-V_o_). The known log of the molecular weights of protein standards were graphed vs their calculated K_av_ values and fit to a logarithmic function (y=m*ln(x)+b).

### FRET-based ssDNA Binding Assays

The FRET-based ssDNA binding assays were performed as described in^29–31,37,54^. Binding to the Cy3-dT60-Cy5 substrate (10 nM) results in a change in FRET between the Cy3 and Cy5 dyes and an observed stoichiometric binding isotherm. RAD51 binding site size was determined by multiplying 10 nM of our Cy3-dT60-Cy5 substrate by the total number of nucleotides to obtain 600 nM nucleotides. Then we divided 600 nM nucleotides by the point at which the two lines meet (200 nM) to obtain a stoichiometry of 3. This means that a RAD51 protein binds to 3 nucleotides of ssDNA as expected.

### Nuclear Extract Preparation

We utilized parental and SWSAP1 knock-out RPE-1 hTERT-immortalized retinal epithelial cells (ATCC CRL-4000) cells ^6^. The day before transfection, approximately 4 million cells were seeded/100 mm in media containing Gibco DMEM (11965-092), 10% FBS (VWR Seradigm 1500-500), % Pen/Strep, and Puromycin. The cells were transiently co-transfected using Lipofectamine 2000 (Invitrogen P/N 52758) and 3 ug of plasmids pHNT-SWSAP1 and pCDNA3-HA-SWS1 in Optimem Mix (Gibco 31985-062) for 5 hrs. Then, the cells were washed 2 x PBS, and phenol red free Gibco DMEM media (A14430-01) containing 100 nM JF-503 dye was added back to the cells in the dark and incubated for 1 hr as described ^63^. Cell lines were maintained in 5% CO_2_ at 37°C.Then in the dark, cells were washed with PBS and pelleted for nuclear extraction (Abcam Ab113474) on ice. Nuclear extracts were aliquoted and snap frozen at −80°C for storage. Once nuclear extracts were obtained, SDS-Page gels were run (without Coomassie dye) and imaged at the appropriate wavelength (488 nm) on the Typhoon Imager (GE). MW bands at approximately 33.6 kDa were observed for the HaloTag-JF503-SWSAP1 protein. The HaloTag-JF503-SWSAP1 was quantified by using recombinant HALO-GFP protein standard (Promega) pre-incubated with JF503 at a 1:1 molar ratio and titrated into single molecule buffer (20 mM Tris pH7.5, 100 mM NaCl, 2 mM ATP, 5 mM MgCl_2_, 5 mM CaCl_2_, 10 mM DTT). Using the HALO-GFP-JF503 standard, protein quantification was determined for each extract and typical yields ranged from 2-10 nM in 30 μL. C-trap data is a summation of >4 nuclear extract preparations.

### Single-molecule measurements

Single-molecule optical tweezer experiments were performed on a LUMICKS C-trap. The LUMICKS C-trap is a confocal trapping system with microfluidics that enables confocal scanning using three excitation lasers (488, 561, and 638 nm) with three emission filters 500-550, 575-625, and 650-750 nm. The concentrations of proteins assayed were: RAD51 80 nM, AlexaFluor488-RPA protein 10 nM, SWSAP1-SWS1 15 nM, and HaloTag-JF503-SWSAP1 (2-30 nM). For kymograph imaging, all proteins were diluted into buffer containing (20 mM Tris pH7.5, 100 mM NaCl, 2 mM ATP, 5 mM MgCl_2_, 5 mM CaCl_2_, 10 mM DTT, and 1 mM Trolox-quinone).

All buffers were degassed and filtered using 0.02 micron filters. The flow cell surface was passivated using BSA (0.1% final) and Pluronics (0.5% final) in PBS buffer. After optical trapping of 4.88 micron streptavidin beads, ssDNA was generated with 10 pN force as described^41,42^. For visualizing RAD51, the ssDNA tethered between the two beads was dipped into the channel containing Cy3-RAD51 protein (80nM) and HaloTag-JF503-SWSAP1 protein (2-10 nM) for 10 seconds and then moved to channel 3 in the buffer listed above with low salt (50 nM). Confocal imaging along the ssDNA was performed in the center of the flow cell and kymographs were collected. For visualizing RPA, the ssDNA tethered between the two beads was dipped into the channel containing AlexaFluor-488-RPA protein (10 nM), moved into channel 3 in the buffer listed above with low salt (50 nM) imaged, and then moved into channel 5 containing non-fluorescent SWSAP1-SWS1 proteins for imaging. Confocal imaging along the ssDNA was performed in the center of the flow cell and kymographs were collected. All experiments were performed with 20% trapping laser overall power, 5% green laser, and 10% blue laser power. Imaging parameters were as follows: 500 ms line time, 100 nm pixel size with a resolution of 300 nucleotides/100 nm. Under these conditions and frame rates, we did not observe bleed through between the green and blue channels.

Importantly, RPA data were collected only with the blue laser. All data are reported as a function of position along the y axis and time on the x axis. Importantly, Cy3-RAD51 photobleaching times at this laser power and imaging rate is ∼616 seconds or 10.26 minutes. Kymographs photobleaching analysis, positional analysis, and colocalization analysis were performed and analyzed using the using Jupyter Notebook and scripts generated by Dr. Matthew Schaich and Dr. David Rueda’s laboratory (Lumicks Harbor)^34^. Contrast was adjusted in kymograph images in the figures. Mean squared displacements (MSDs) were used to track particle motion and individual tracked filaments were used to calculate MSDs as described in^34^. Diffusivity was calculated based on a linear curve of pure diffusion by performing ordinary least squares, an unweighted fit of the data, of the estimated MSD values. Data were plotted using GraphPad Prism.

### D-loop Assays

These assays were performed using RPA-coated ssDNA which would be the physiologically relevant species, removes any secondary structure within ssDNA, and binds displaced strands within the reaction. We also assayed two different constructs of the human Shu components. The first was a GST-SWS1-SWSAP1 recombinant protein and the second was a cleaved SWSAP1-SWS1 without any large recombinant tags. To do this we incubated 3 μM of 90 mer ssDNA labeled with FITC which is complementary to a sequence in the pBluescript KG plasmid with RPA at 100 nM^64^. Then we challenged this complex with RAD51 (1 μM) or RAD51-SWSAP1-GST-SWS1 complexes at increasing concentrations (RAD51 250 nM: SWSAP1-GST-SWS1 5 nM, 10 nM, 20 nM, 40 nM, 80 nM, 160 nM, 360 nM and 1 μM) for 5 min at 37°C. We initiated the reaction by adding pBluescript KG plasmid at 50 μM bp concentration for 15 min at 37°C, stopped the reaction by incubation with SDS and Proteinase K, and ran the reaction products on an agarose gel. D-loops were visualized on a Typhoon instrument (GE) by imaging and exciting at Cy2 (495 nm). Then, % D-loops were quantified using the Typhoon software (GE) and normalized to RPA-RAD51 only reactions. Experiments were also performed with untagged SWSAP1-SWS1 where the GST was cleaved.

### Yeast-two-hybrid (Y2H) Assay

Yeast-two-hybrid (Y2H) plasmids were constructed from the GAL4 DNA activating domain (pGAD) and GAL4 DNA-binding domain (pGBD). pGBD-SWSAP1 and pGAD-SWS1 were previously published ^7^.The 16 selected cancer-associated SWSAP1 mutants were made through pGBD-SWSAP1 site-directed mutagenesis. The pGAD-SWS1 and pGBD-SWSAP1 plasmids were co-transformed into the *S. cerevisiae* PJ69-4a strain and plated on media selecting for yeast transformants expressing the two plasmids (SC-LEU-TRP) and grown for 48 hours at 30°. For the Y2H assay, 3-4 individual colonies transformed with the according plasmids were selected and cultured overnight in selective media (SC-LEU-TRP). The cultures were then diluted to OD_600_ 0.5 and grown to early log phase (4 hrs). They then were diluted to OD_600_ 0.2 and 5 μL of the culture was spotted onto medium selecting for the plasmids (SC-LEU-TRP), medium selecting for the expression of the HIS3 reporter gene (SC-LEU-TRP-HIS), or medium selecting for the HIS3 reporter and stressing the strength of the interaction through the inclusion of a competitive inhibitor (SC-LEU-TRP-HIS+3AT). Plates were incubated for 3 days at 30°C and photographed after 48 and 72 hours. The experiments were done in triplicate. All Y2H images were adjusted identically for brightness and contrast using Adobe Photoshop.

### Western Blotting of Y2H Assays

Yeast expressing the constructed pGAD and pGBD constructs were grown overnight in 3ml of YPD. Whole-cell lysates of equal cell numbers (0.75 OD_600_) were extracted by TCA precipitation. Protein was separated on 4-15% gradient gels and transferred to PVDF membranes. The SWSAP1 expression was detected with SWSAP1 antibody (1: 500 dilution; ab185360), the loading control was detected using a Kar2 antibody (1:2000), and IR dye secondary antibodies from LiCor Biosciences (1:1000 dilution). Protein was detected on a LiCOr CLX scanner and brightness, and contrast was adjusted using the Imager Software.

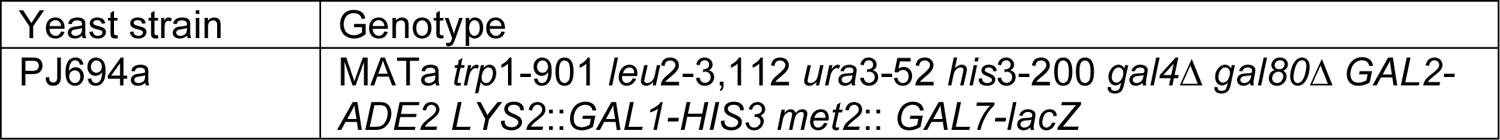

### Native PAGE Complex Formation

Human SWSAP1-SWS1 (4 & 6 μM) and RAD51 (7.5 μM) were incubated in a final volume of 20 μL for 10 min at RT in Native PAGE Sample Buffer (4x) (50 mM Bis Tris pH 7.2, 50 mM NaCl, 50% Glycerol, 0.001% Ponceau S (Invitrogen)). Then, 0.5 μL of Native PAGE Sample G-250 Sample Additive (Invitrogen) was added to bind protein complexes for 5 min at 25 °C. Samples were then loaded and run on 4-16% native PAGE gel (Invitrogen) in 0.5% TBE buffer pH 8.5 at 4 °C for 3 hrs. The native PAGE gel was stained with Coomassie, destained, and shifted bands were cut out for protein composition analysis using mass spectrometry. Excised bands corresponded to protein complexes of RAD51-SWSAP1-SWS1. Peptides for each corresponding proteins were identified in each sample.

### Clonogenic Survival Assays

Clonogenic survival assays of the parental RPE-1, and two independent CRISPR/Cas9 clones of *SWS1* (*sgSWS1-C1* and *sgSWS1-C2*) and *SWSAP1* (*sgSWSAP1-C1* and *sgSWSAP1-C2*) were performed^6^. The indicated cell lines were seeded into 6 well plates and then trypsinized and cell number counted using a Z Coulter Counter (Beckman). Approximately 200 cells were seeded into 60 mm dishes and exposed 25, 50, or 100 µM of APE1 inhibitor (APE1 inhibitor III; Calbiochem) or 0, 1, 2, 5, 10, 25 µM Olaparib for approximately 8-14 days. The cells were stained with crystal violet by first rinsing the cells with PBS twice, then fixing them in methanol for 20 min., followed by crystal violet (0.5% crystal violet and 20% methanol) for 30 min. For APE1 and Olaparib inhibition, the colonies were counted and compared to the untreated control to calculate the relative survival or colony area and intensity using the ColonyArea ImageJ plugin. All cellular assays were performed in biological triplicate with different passages of both clones from each genetic knockout. The average relative survival for each clone was analyzed two-way ANOVA by comparison to the corresponding RPE-1 treated cell line.

### Author contributions

S.R.H. & K.A.B. designed the biochemical, biophysical, and cellular studies. S.R.H. expressed and purified all recombinant proteins, labeled all Cy3- and Cy5-RAD51 filaments, performed FRET-based assays, mass photometry assays, Olaparib survival assays, native PAGE analysis, and SDS-Page analysis of recombinant proteins. K.O. performed yeast-two hybrid and western blot analysis. C.S. performed yeast-two-hybrid analysis. M.A.S. provided C-trap training, and aid with data analysis of single-molecule kymograph via shared scripts (Now on harbor). J.M. performed the clonogenic survival assays with APE1 inhibition. S.R.H and H.L.R performed the clonogenic survival assays with Olaparib. K.D. performed western blots of SWSAP1 from Y2H experiments. M.S. provided protocols for FRET-based RAD51 based assays and pCH1-RAD51 plasmid, and S.R.H. performed mass photometry experiments in M.S. laboratory. S.R.H. performed single-molecule experiments in B.V.H. laboratory using the Lumicks C-trap Instrument.

## Acknowledgements

The authors would like to thank Dr. Luke Lavis for generously sharing the Janelia Farm HaloTag dyes. The authors would like to thank Artur Karczmarczyk from David Rueda’s lab for helping modulate their scripts to enable our blue and green data analysis. The authors would like to thank Dr. Shixin Liu (Rockefeller University) for providing the AlexaFluor488-RPA protein for the use in these studies.

## Materials & Correspondence

Kara Bernstein and Sarah Hengel can be contacted for correspondence and material requests at.

## Data Availability

Authors can confirm that all relevant data are included in the published article in addition to supplementary information files.

