## SupplementFigures for "The human Shu complex promotes RAD51 activity by modulating RPA dynamics on ssDNA"

### Supplementary Figure 1 Hengel et al., 2024

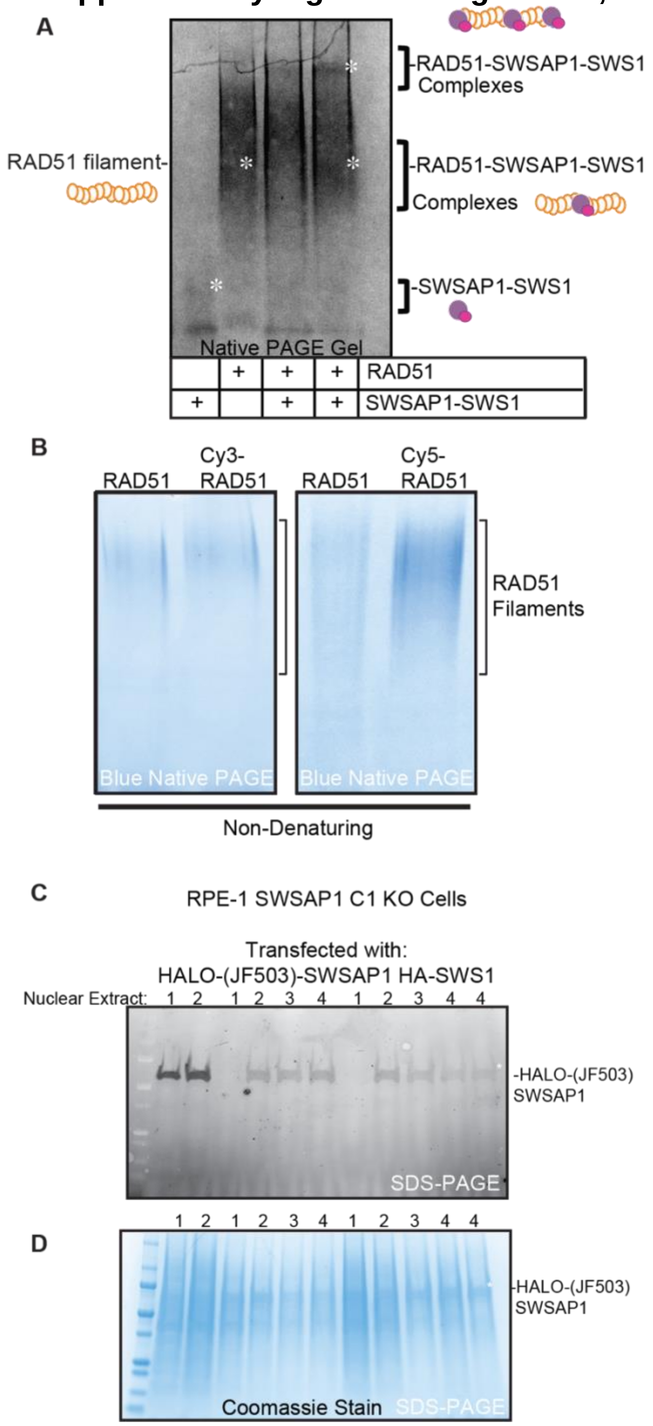


**Supplementary Figure 1: SWSAP1-SWS1 interact with RAD51 and Cy3 or Cy5-labeled RAD51 is functional.** A) Native PAGE gel was run in the presence of $MgCl_2$ in reactions containing SWSAP1-SWS 4 μM (well 1), RAD51 7.5 μM (well 2), SWSAP1-SWS1 4 μM and RAD51 7.5 μM (well 3), and SWSAP1-SWS1 6 μM and RAD51 7.5 μM (well 4). The gel was stained with Coomassie blue and the indicated samples (white *) were sent for mass spectrometry peptide analysis. B) RAD51, Cy3-RAD51, and Cy5-

RAD51 (2 μM final) were loaded onto Native PAGE gels, run for 4 hours and stained with Coomassie blue before imaging. C) Nuclear extracts isolated from *sgSWSAP1* RPE-1 knock out cells transiently transfected with HaloTag-SWSAP1 and HA-SWS1 were run on SDS-PAGE gels and incubated with JF503. The numbers at the top of the gel correspond to different cell passages. The protein was visualized by imaging on a Typhoon GE Imager by exciting at 488 nm. D) Nuclear extracts in (C) were run on an SDS-PAGE gel and stained with Coomassie blue. The HaloTag-JF503-SWSAP1 protein is indicated to the right of the image.

### Supplementary Figure 2 Hengel et al., 2024

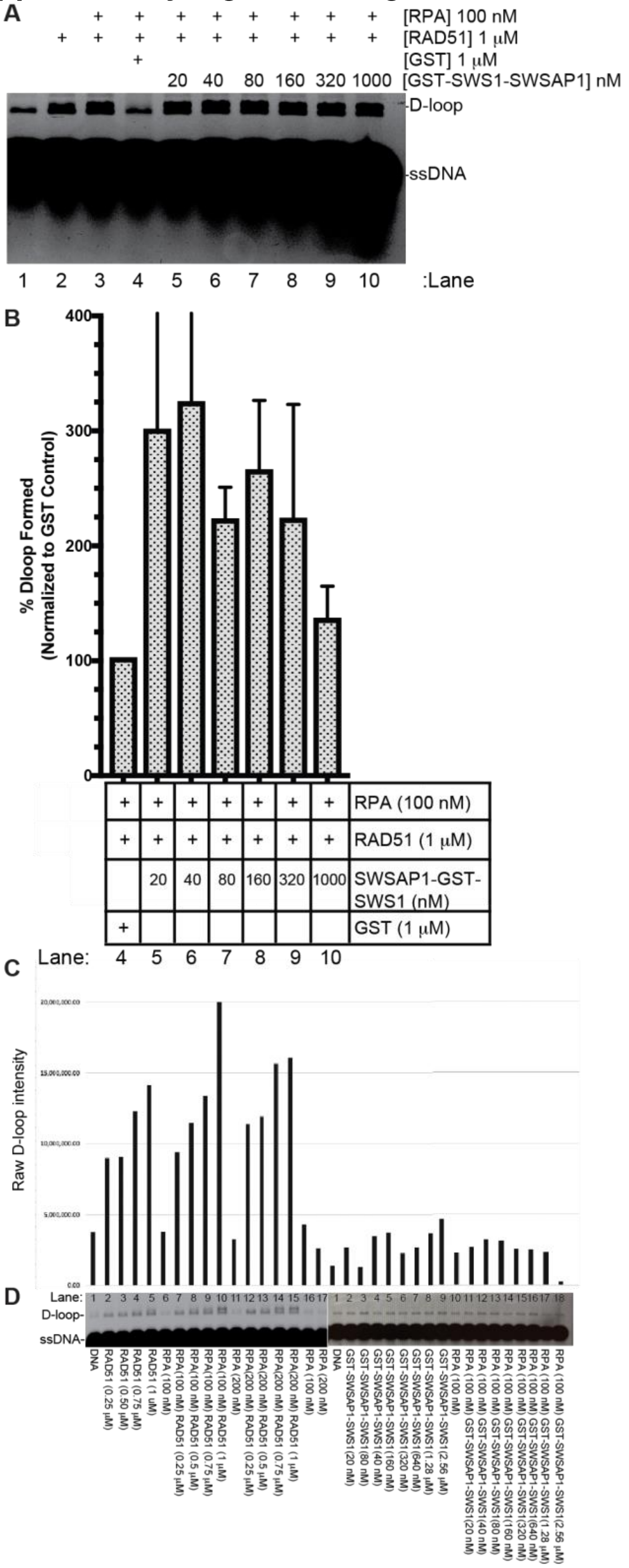


**Supplementary Figure 2: GST-SWSAP1-SWS1 stimulates RAD51-dependent D-loop formation on RPA-coated ssDNA.** A) Representative agarose gel of D-loop products. B) Quantifications of D-loop reactions show a 2x stimulation of D-loop product formed even at the lowest SWSAP1-GST-SWS1 protein concentration (5 nM). Experiments were performed in triplicate and standard deviations are shown. C) Quantification of D-loop reactions shown in (D). D) Representative agarose gel of D-loop products for RAD51, RPA and RAD51, GST-SWSAP1-SWS1, and RPA with GST-SWSAP1-SWS1 at the indicated concentrations. Note that the right quantification graph was cut from lines 9-18 to better align with the lanes.

**Supplementary Figure 3 Hengel et al., 2024**

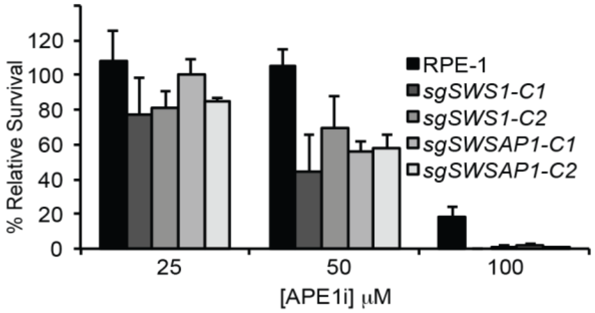


**Supplementary Figure 3: SWS1 and SWSAP1 CRISPR/Cas9 knockout RPE-1 cells are sensitive to APE1 inhibition.** Clonogenic survival assays of the parental RPE-1, and two independent CRISPR/Cas9 clones of *SWS1* (*sgSWS1-C1* and *sgSWS1-C2*) and *SWSAP1* (*sgSWSAP1-C1* and *sgSWSAP1-C2*). All cellular assays were performed in biological triplicate with different passages of both clones from each genetic knockout. The indicated cell lines were exposed to the indicated concentration of APE1 inhibitor (APE1 inhibitor III; Calbio) for approximately 14 days and then stained with crystal violet. The colonies were counted and compared to the untreated control to calculate the relative survival and standard deviations are plotted. The average relative survival for each clone was analyzed two-way ANOVA by comparison to the corresponding RPE-1 treated cell line (SWS1-C1 $p = 0.002$, SWS1-C2 $p = 0.002$, SWSAP1-C1 $p = 0.0497$, and SWSAP1-C2 $p = 0.0145$).
